# An L-threonine-inducible biosensor derived from the tryptophan-repressed *trp* promoter of *Escherichia coli* for high-throughput screening of L-threonine overproducers

**DOI:** 10.64898/2026.10.01.755634

**Authors:** Buli Su, Peixuan Lai, Xiong Xiong, Yonghong Wang, Man Li, Honghui Zhu, Ming-Rong Deng

## Abstract

L-Threonine is an essential amino acid widely used in the feed, food, and pharmaceutical industries, and microbial fermentation is now the dominant route for its production. Further increases in producer titers depend on random mutagenesis coupled with high-throughput screening (HTS), but progress has been limited by the lack of a biosensor with a broad operating range, high sensitivity, and target specificity. Here, we show that L-threonine induces the tryptophan-repressed *trp* promoter of *Escherichia coli*, a response that, to our knowledge, has not been described previously. Deletion analysis showed that both the *trpO* operator and the *trpL* attenuation element contribute to induction, and that the lowest basal expression occurs only when both elements are present. By optimizing the ribosome-binding site, the promoter –10 region, and *TrpR* expression, we constructed an L-threonine biosensor on a pACYC backbone. The sensor responded linearly to L-threonine from 0.1 to 30 g/L (R² = 0.9895), with an induction ratio of up to 51.7-fold. Among the 20 proteinogenic amino acids tested, only L-threonine produced a detectable response. The biosensor distinguished strains with different L-threonine production capacities and guided the directed evolution of homoserine kinase (ThrB) at a conserved motif, yielding variants with brighter reporter fluorescence and increased L-threonine production. This work provides a high-performance biosensor based on native *E. coli* regulatory elements for screening L-threonine overproducers.

## 1. Introduction

L-Threonine belongs to the aspartate family of amino acids and is essential in animal nutrition, where it is often one of the growth-limiting amino acids in corn-soybean meal-based diets (Alabi and Adedokun 2025). Demand has grown with the feed, food, and pharmaceutical industries, and microbial fermentation now supplies most of the market. Over the past two decades, engineered *Escherichia coli* strains have reached titers above 170 g/L in fed-batch cultures (Zhao et al. 2024). Much of this progress has resulted from rational rewiring of biosynthetic flux, but the pathway is tightly regulated by feedback mechanisms (Wang et al. 2023; Zhao et al. 2025a), limiting further gains from these well-characterized components. Additional improvement therefore depends largely relies on random or semi-rational mutagenesis, whose value is determined by how efficiently rare beneficial mutants can be recovered from large libraries.

Genetically encoded biosensors have emerged as a general solution to this screening bottleneck (Shi et al. 2018; Yu et al. 2023). By coupling the metabolite concentration to fluorescent-reporter, these sensors convert intracellular production into a signal that can be measured by flow cytometry or plate readers. They have enabled high-throughput screening for several amino acids. Successful examples use diverse sensing elements, including an L-lysine-inducible promoter, a cysteine-responsive transcription factor, an L-tryptophan biosensor, and glycine-responsive riboswitch– and transcription-factor-based elements (Gao et al. 2022; Hong et al. 2022; Li et al. 2025; Peng et al. 2026). In another design, an engineered isoleucyl-tRNA synthetase couples isoleucine recognition directly to cell growth (Sun et al. 2021). Because each amino acid requires its own sensing element, a sensor that performs well for one target cannot be assumed to work for another.

For L-threonine, a sensing element with comparable performance has not yet emerged. The first threonine-responsive promoters were identified by comparing the proteomes *E. coli* grown with and without the amino acid. One of them, the fusion promoter *cysJHp*, enabled screening of threonine overproducers, although its approximately fourfold signal span limited quantitative resolution (Liu et al. 2015). A fast coupled-enzyme assay provided an alternative microtiter-format method for quantifying L-threonine without a genetically encoded reporter (Liu et al. 2014). A dual-response circuit combining the threonine-sensing riboswitch *thrL* with a *lacI*/P*_trc_* amplification module reached a 4.2-fold dynamic range over 0-20 g/L and was used successfully to evolve thrA (Su et al. 2024). A CysB-based sensor responding to threonine up to approximately 4 g/L, with a 5.6-fold signal increase, was also reported (Zhao et al. 2025b), and the mutated regulator SerR^F104I^ recognized both threonine and proline (Pu et al. 2025). These devices demonstrate the feasibility of threonine sensing, but each has limitations. The promoter– and regulator-based sensors have narrow operating windows relative to the titers achieved by modern producers, and provide modest signal amplitudes. The riboswitch-based circuit, in contrast, requires complex signal amplification. A single sensor combining a broad linear range, high sensitivity, and specificity for the L-threonine has been lacking.

The tryptophan (*trp*) operon of *E. coli* is one of the best-characterized regulatory systems in bacteria. When bound to tryptophan, the repressor TrpR blocks transcription initiation at the *trpO* operator, which overlaps the promoter –10 region, whereas the leader peptide *trpL* mediates transcriptional attenuation in response to tryptophan availability (Li et al. 2025; Yanofsky 2003; Yanofsky et al. 1981). These mechanisms silence the operon when tryptophan is abundant, so the promoter is generally regarded as a strictly tryptophan-repressed element. During a transcriptomic and proteomic study of a post-transcriptional stimulatory effect of L-threonine on recombinant protein expression, we observed marked upregulation of the *trpEDCBA* cluster. This finding was unexpected, because tryptophan biosynthetic genes are not known to respond to threonine. It prompted us to ask whether L-threonine can induce a promoter normally silenced by its cognate, tryptophan.

Here, we report that L-threonine induces the tryptophan-repressed *trpEDCBA* promoter of *E. coli*, a response that, to our knowledge, has not been reported previously. The *trpO* operator and the *trpL* attenuation element together produced the tightest OFF state, so we used both elements to convert the native promoter into a biosensor. We optimized the ribosome-binding site, the promoter –10 region, plasmid copy number, and TrpR abundance. The resulting biosensor responded linearly to L-threonine over 0.1-30 g/L, was specific for L-threonine among the 20 amino acids tested, discriminated strains with different production capacities, and supported directed evolution of homoserine kinase (ThrB).

## 2. Materials and methods

### 2.1 Strains, plasmids and culture conditions

*Escherichia coli* Turbo and W3110 were used as the hosts for biosensor construction and characterization, respectively. Two L-threonine-producing strains with different biosynthetic capacities were used for functional validation. All strains, plasmids, and primers used in this work are listed in Tables S1 and S2. Cells were routinely grown in Luria-Bertani (LB) medium supplemented with 5 g/L MOPS at 37 °C. Kanamycin and other antibiotics were added as required to maintain plasmids. Liquid cultures were shaken at 200 rpm in 150-mL flasks containing 30 mL of medium.

### 2.2 Construction of biosensor plasmids

The promoter region of the *trpEDCBA* operon, including the *trpO* operator and the *trpL* leader sequence, was amplified from the *E. coli* W3110 genome and fused to the *egfp* coding sequence. Plasmids were assembled with the MultiF Seamless Assembly Mix (ABclonal Technology Co., Ltd., China) according to the manufacturer’s instructions. PCR and colony PCR were performed with Phanta Flash DNA polymerase and Green Taq Mix (Vazyme Biotech Co., Ltd., China), respectively. The native ribosome-binding site (RBS) upstream of *egfp* was replaced with a strong synthetic RBS sequence (from pET30a). For the deletion analysis, *trpR* was deleted from the chromosome using CRISPR-Cas9 system, whereas trpL deleted from the biosensor plasmid by the whole-plasmid PCR. To examine the contribution of individual regulatory elements, the synthetic constitutive promoter P*_J23119_* was combined with *trpO*, *trpL*, or both, and the resulting variants were placed upstream of *egfp*.

Promoter –10-region mutants were constructed by site-directed mutagenesis of the native *trp* promoter. The –10 hexamer was changed from TTAACT to TATAAT (the three-position mutant, –10A) or to TATACT (two-position mutant, –10B). Plasmid backbones with different copy numbers, namely pCDF (CloDF13 ori), pET30a (pBR322 ori), pSC101 (SC101 ori), and pACYC (p15A ori), were tested as biosensor carriers. To tune TrpR abundance, trpR was expressed from its native promoter or from constitutive promoters of different strengths (P*_J23119_*, P*_J23104_*, P*_J23110,_* and P*_trpR_*), cloned on the same or a compatible plasmid.

### 2.3 Cultivation and fluorescence measurement

Strains harboring biosensor plasmids were precultured in LB medium at 37°C for 12 h and then inoculated into fresh medium to an initial OD_600_ of approximately 0.05. L-threonine was added to the indicated final concentrations, and cultures were grown under the same conditions for 24 h. Cell density (OD*_600_*) and eGFP fluorescence were measured with a Thermo Scientific Varioskan LUX) microplate reader using 488-nm excitation and 509-nm emission. Specific fluorescence was calculated as the ratio of fluorescence intensity to OD*_600_* (eGFP/OD*_600_*).

For the specificity test, each of 20 amino acids was added at 20 mM, and the response was compared with that obtained with L-threonine. To exclude interference from impurities, L-threonine from three different commercial suppliers (Sigma-Aldrich, Macklin, and Aladdin) was assayed under identical conditions.

### 2.4 Functional validation in L-threonine-producing strains

The optimized biosensor plasmid was introduced into two strains with different L-threonine production capacities. Transformants were cultivated in LBGM fermentation medium, consisting of LB supplemented with 5 g/L glycerol and 5 g/L MOPS. Specific fluorescence and extracellular L-threonine concentration were measured at the end of cultivation. The relationship between the fluorescence output and the measured titer was used to determine whether the sensor could discriminate strains with different productivities.

### 2.5 Saturation mutagenesis of thrB and biosensor-based screening

The conserved A12 motif of homoserine kinase (ThrB) was selected based on sequence alignment of ThrB orthologs (Petit et al. 2018). Saturation mutagenesis at A12 was performed with primers carrying an NNK degenerate codon. The thrB coding sequence was cloned into the pET30a vector under the control of the P*_J23104_* promoter, and the mutant library was constructed by DpnI digestion and transformation. The library was introduced into WS000 cells harboring the pACYC-based biosensor, and transformants were spread on LBGM plates. Colonies with elevated fluorescence were picked and cultivated in LBGM medium, and their L-threonine production was quantified. Mutations were confirmed by DNA sequencing.

### 2.6 L-threonine quantification

L-Threonine concentrations were determined by high-performance liquid chromatography (HPLC) on an Agilent 1260 system equipped with a Thermo ODS-2 Hypersil C18 column, using o-phthalaldehyde (OPA) pre-column derivatization, as described previously (Su et al. 2024).

### 2.7 Statistical analysis

All experiments were performed with at least three biological replicates, defined as independent cultures inoculated from separate colonies. Data are presented as means ± standard deviations. Statistical significance was evaluated with two-tailed Student’s t-tests, with *p* < 0.05 considered significant.

## 3. Results and discussion

### 3.1 L-Threonine induces the tryptophan-repressed *trp* promoter of *E. coli*

This work began with a serendipitous observation made while we were investigating the post-transcriptional stimulatory effect of L-threonine on recombinant protein expression (Su et al. 2024). The RNA-seq and proteomic data were deposited in the Genome Sequence Archive (GSA) server and OMIX at the Big Data Center (https://ngdc.cncb.ac.cn; GSA accession No. CRA050252; OMIX ID. OMIX020827). Transcriptomic and proteomic comparisons showed that the *trpEDCBA* genes were markedly upregulated in cells grown with L-threonine (Fig. 1A). This finding was unexpected for two reasons. The *trp* operon is among the most tightly regulated units in *E. coli*, being silenced by the TrpR repressor at the *trpO* operator and by transcriptional attenuation at the *trpL* leader; both mechanisms respond to tryptophan. Threonine, in contrast, had not previously been linked to the regulation of tryptophan biosynthetic genes. We therefore tested whether the promoter itself responds to L-threonine by fusing the *trpEDCBA* promoter region to an *egfp* reporter and monitoring fluorescence in *E. coli* W3110 (Fig. 1B).

**Fig. 1.**
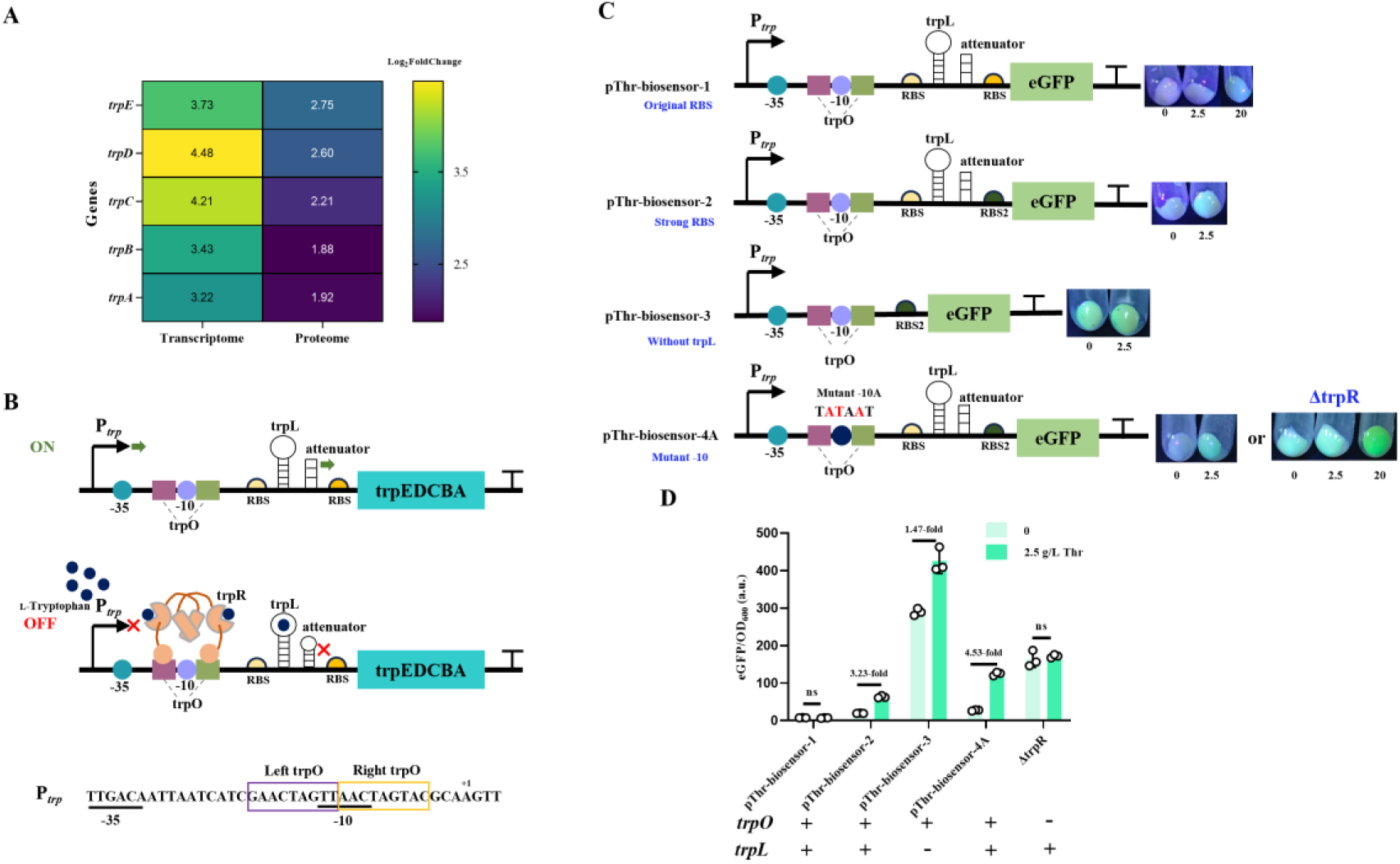
The tryptophan-repressed *trp* promoter of *E. coli* responds to L-threonine. (A) Transcriptomic and proteomic data from *E. coli* grown in LB medium containing 2.5 g/L L-threonine; the *trpEDCBA* gene cluster is indicated. (B) Schematic of the *trpEDCBA* promoter (P*_trp_*), showing the *trpO* operator and the *trpL* leader. TrpR is a dimer, in the absence of tryptophan, the promoter is in an open (transcriptionally permissive) state; in the presence of tryptophan, tryptophan-bound TrpR acts at *trpO* and at *trpL*, and the promoter is closed. *trpO* overlaps the –10 region of P*_trp_*. (C) Schematics of the biosensor constructs and representative cell images, P*_trp_* with its native ribosome binding site (RBS) (*trpO* and *trpL* present); P*_trp_* with a strong synthetic RBS (*trpO* and *trpL* present); Ptrp with trpL deleted; P*trp* with trpR deleted (eliminating the effect of trpO). (D) Specific fluorescence (eGFP/OD*_600_*) and signal-to-background ratio of the constructs in (C) under uninduced conditions and after induction with 2.5 g/L L-threonine. ns: no significant difference by Student’s t-test.

In the initial construct, which retained the native ribosome-binding site (RBS) of the operon, no induction was detected when L-threonine was added at 2.5 or 5 g/L (Fig. 1C and Fig. S1A). The promoter responded when the threonine concentration reached 20 g/L (Fig. S1A), indicating that L-threonine can derepress the *trp* promoter but that the native configuration produces only a weak signal. Replacing the native RBS with a stronger sequence changed this behavior: the reporter was induced even at low threonine concentrations, with a signal-to-background ratio of 3.23-fold (Fig. 1D).

To determine which of the two known regulatory circuits was involved, we deleted trpR to remove repression and *trpL* to remove attenuation. Deletion of *trpR* increased basal expression, yet the sensor was induced only at high threonine concentrations (Fig. S1C). Deletion of *trpL* increased the basal expression further, and the response to L-threonine remained detectable, but the signal-to-background ratio was only 1.47-fold (Fig. 1D). These elements therefore act together: neither one alone reproduced the tight OFF state and the strong threonine-dependent response of the intact promoter. Only when *trpO* and *trpL* were both present was basal expression lowest while a strong threonine-dependent response was retained.

We next asked whether this behavior could be reproduced with a defined synthetic promoter. When the constitutive promoter P*_J23119_* was combined with *trpO*, *trpL*, or both, the variants carrying complete set of regulatory elements again showed the lowest background and the clearest threonine-dependent induction, matching the native promoter (Fig. 2A). As in the native promoter, *trpO* and *trpL* together made the synthetic promoter responsive to L-threonine, whereas neither element alone produced the tightest OFF state (Fig. 2B). The molecular mechanism by which L-threonine elicits this response remains unknown. Tryptophan analogues lacking the α-amino group, such as indole-3-propionic acid, are established inducers of the *trp* operon (Hyde et al. 1991; Lawson and Sigler 1988). Crystal structures show that, without the amino group, the ligand reorients so that its carboxyl group faces the DNA-binding surface and repels the operator backbone, preventing operator binding. By analogy, L-threonine may derepress the promoter through a related perturbation of tryptophan-responsive regulation. This finding extends the class of molecules known to derepress the *trp* operon from tryptophan analogues to a proteinogenic amino acid.

**Fig. 2.**
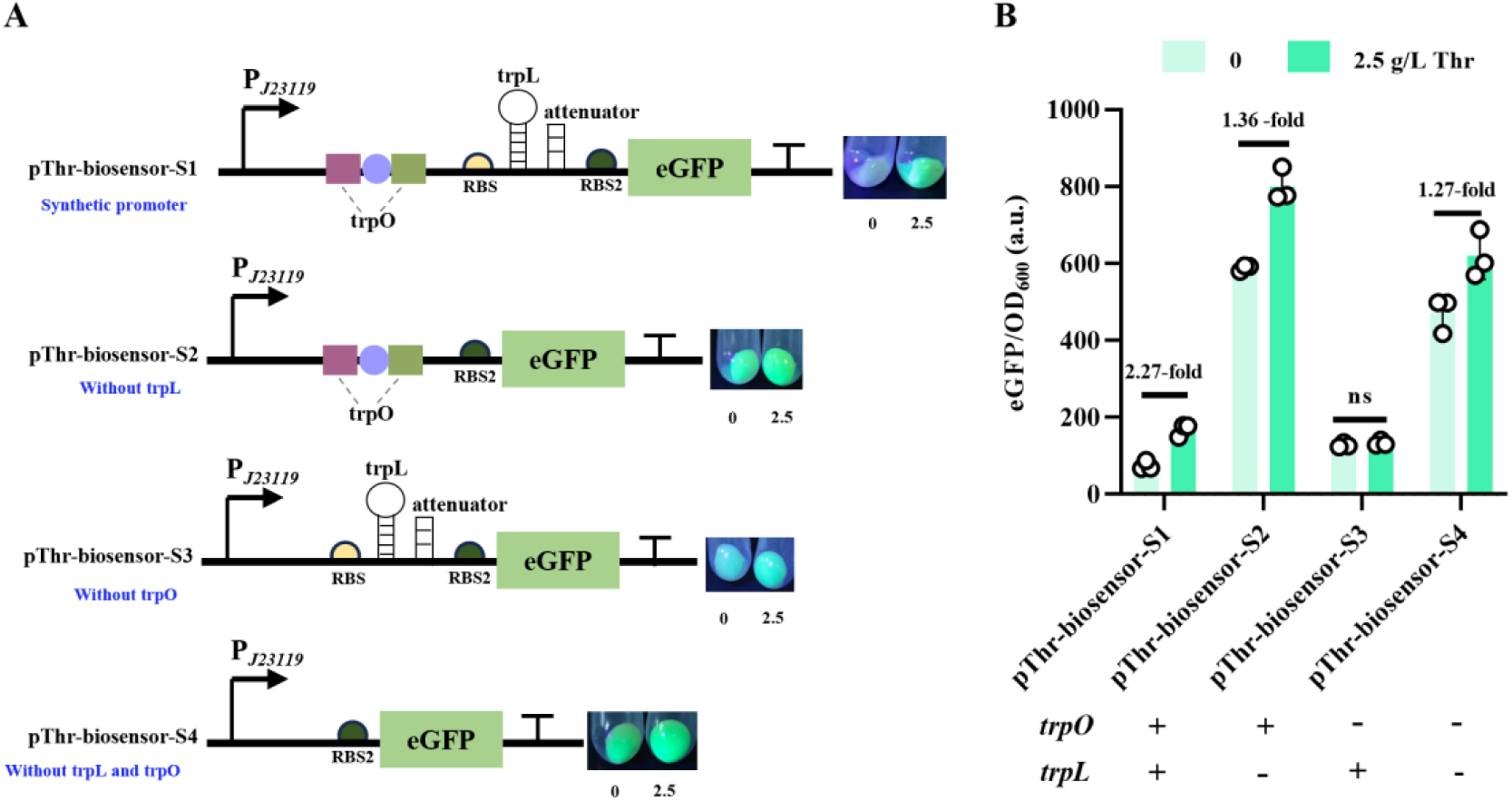
A synthetic promoter reproduces the threonine response of the native *trp* **regulatory elements.** (A) Schematics and representative cell images of sensors based on the constitutive promoter P*_J23119_*; P*_J23119_* carrying *trpO* and *trpL*; P*_J23119_* with *trpL* deleted; P*_J23119_* with *trpO* deleted; and P*_J23119_* without *trpO* or *trpL*. (B) Specific fluorescence (eGFP/OD*_600_*) and signal-to-background ratio of the constructs in (A) under uninduced conditions and after induction with 2.5 g/L L-threonine.

The ability of the *trp* promoter to serve as the core of an amino acid sensor is not apparent from the existing. TrpR itself can be re-engineered, directed evolution can redirect its DNA-binding specificity (Ellefson et al. 2018), and rational mutation of its ligand-binding pocket can alter ligand preference and dynamic range (Gong et al. 2022). Recent tryptophan-responsive biosensors have instead used RNA-binding attenuation proteins to avoid the complications of repressor tuning (Xu et al. 2026). The finding that the *trp* promoter is derepressed by L-threonine, without re-engineering of TrpR, therefore differs from both strategies. It also suggests that promoter elements regarded as specific to one amino acid may respond to other metabolites.

### 3.2 Stepwise engineering of the promoter and the sensing context improve sensor performance

The native *trp* promoter responded to L-threonine, but its output window was far from ideal, so we set out to improve it. Because the –10 region of the *trp* promoter overlaps the *trpO* operator, changes in this region affect both transcription strength and repressor binding, making it a natural target for engineering (Fig. 1B).

We first mutated three positions in the –10 region, converting the non-consensus sequence TTAACT to the strong consensus sequence TATAAT (designated –10A) (Fig. 1C). This change strengthened the response to L-threonine but also substantially increased uninduced expression, so the overall signal-to-background ratio reached only 4.53-fold (Fig. 1D). The higher background presumably reflected disruption of TrpR binding at the overlapping *trpO* site, which weakened repression. We therefore reduced the number of substitutions, changing the same hexamer to TATACT (designated –10B) (Fig. 3). This variant restored consensus bases at two of the three positions altered in – 10A while leaving the operator-proximal base unchanged. The –10B variant showed a much lower basal fluorescence, close to that of a W3110 strain without the reporter (Fig. S1D), and retained a clear threonine-dependent response, with a signal-to-background ratio of 6.06-fold (Fig. 3). This result illustrates a common biosensor trade-off: strengthening the promoter core raises the induced signal but can weaken the OFF state when the operator and promoter overlap.

**Fig. 3.**
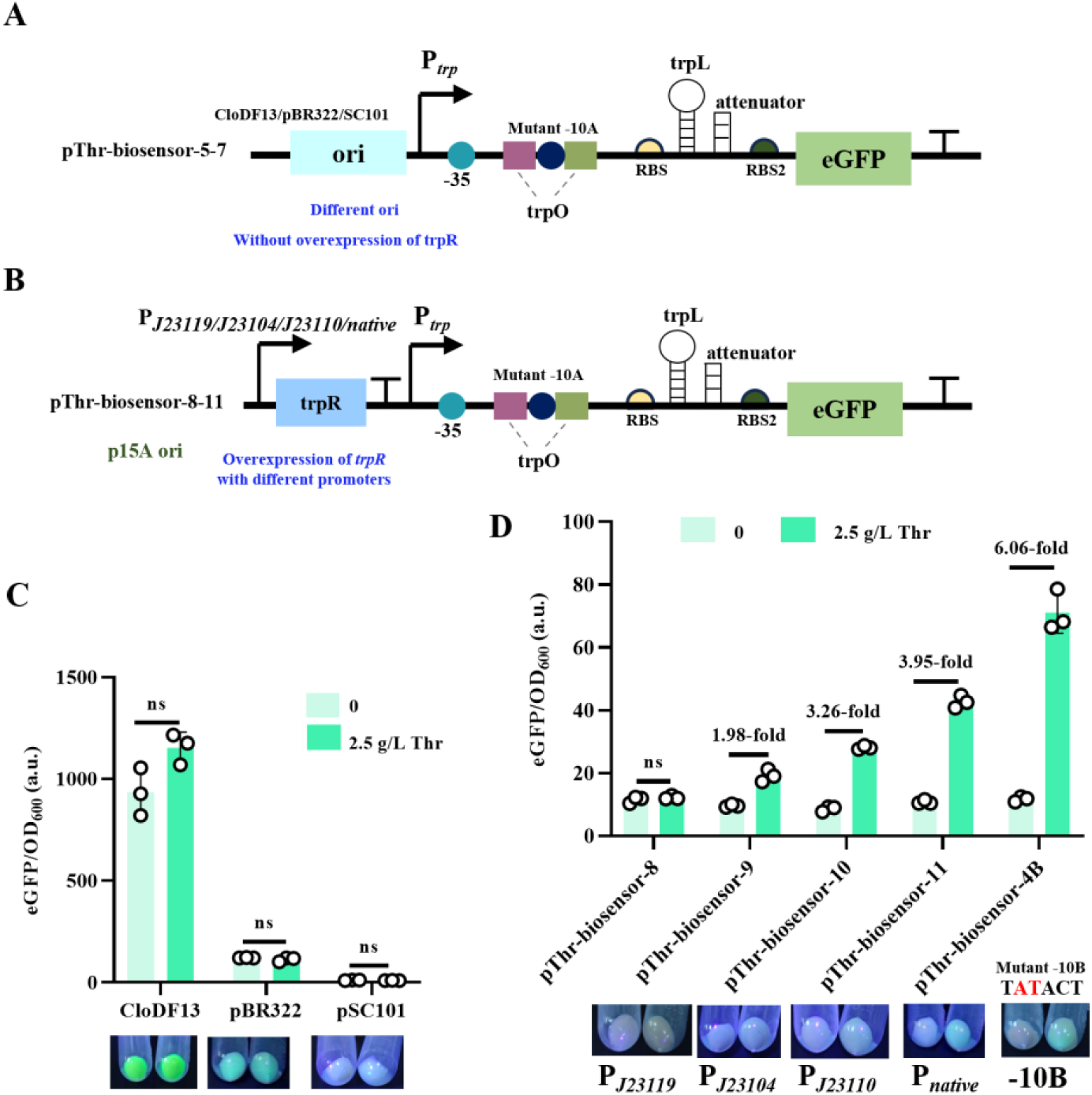
Effects of plasmid copy number and TrpR abundance on the biosensor. (A) Schematics of the sensor on replicons with different copy numbers (CloDF13, pBR322, and SC101). (B) Schematics of the sensor with trpR expressed from promoters of different strengths (P*_J23119_*, P*_J23104_*, P*_J23110_*, and the native trpR promoter). (C) Specific fluorescence and representative cell images of the replicon variants in (A) under uninduced conditions and after induction with 2.5 g/L L-threonine. (D) Specific fluorescence and signal-to-background ratio of the trpR-expression variants and the – 10B biosensor in (B) under uninduced conditions and after induction with 2.5 g/L L-threonine. ns, no significant difference by two-tailed Student’s t-test.

The sensing context surrounding the promoter also affected performance. We compared four plasmid backbones spanning a broad range of copy numbers: pCDF (CloDF13 ori), pET30a (pBR322 ori), pSC101 (SC101 ori), and pACYC (p15A ori) (Fig. 3A). Only pACYC, which has an intermediate-to-low copy number, yielded a biosensor with clear threonine-dependent induction (Fig. 3C). Higher-copy plasmids presumably increase the gene dosage of the promoter and chromosomal trpR enough to disturb the repressor-operator balance, whereas the lowest-copy construct may provide too few reporter molecules to generate a measurable signal. We varied TrpR abundance in parallel by expressing it from promoters of different strengths (P*_J23119_*, P*_J23104_*, P*_J23110_*, and P*_trpR_*) (Fig. 3B). Overexpression shifted induction toward high threonine concentrations and reduced the signal-to-background ratio (Fig. 3D). The sensor therefore performs well only when repressor abundance and the operator-containing promoter are appropriately balanced, a parameter that can be overlooked when a sensor is transferred between plasmid contexts.

The present sensor is unusual because its main tuning element was the operator embedded in the –10 region, rather than the regulator itself. Transcription-factor-based sensors are typically tuned through transcriptional or translational control of the regulator (Zhou and Zhang 2023). Because this sensor reports the intracellular L-threonine pool directly, it could in principle be coupled to dynamic regulation of production pathways as well as used for endpoint screening (Xu et al. 2020). Preserving the operator in the –10B variant was decisive. Although the induced signal was somewhat lower than that of –10A, the background decreased more sharply, improving the signal-to-background ratio. The resulting construct was used for all subsequent characterization. Because the operator and promoter overlap, any change to the promoter core also affects repressor binding. Similar overlaps in other metabolite-repressed promoters may provide a comparable tuning strategy.

### 3.3 The optimized sensor reports L-threonine over a broad linear range and is specific for the target amino acid

We first characterized the –10B sensor in cultures supplemented with L-threonine. As the concentration increased from 0 to 30 g/L, the specific fluorescence of *E. coli* W3110 harboring the biosensor increased approximately linearly. The reporter output fit a straight line with R² = 0.9895. The lowest tested concentration distinguishable from the control was 0.1 g/L, and the response continued to increase up to the highest concentration tested, 30 g/L (Fig. 4). The maximum-to-minimum signal ratio across the tested range was 51.7-fold. To our knowledge, no previously reported L-threonine biosensor combines broad linear range, a low detection threshold, and a large induction amplitude in a single device.

**Fig. 4.**
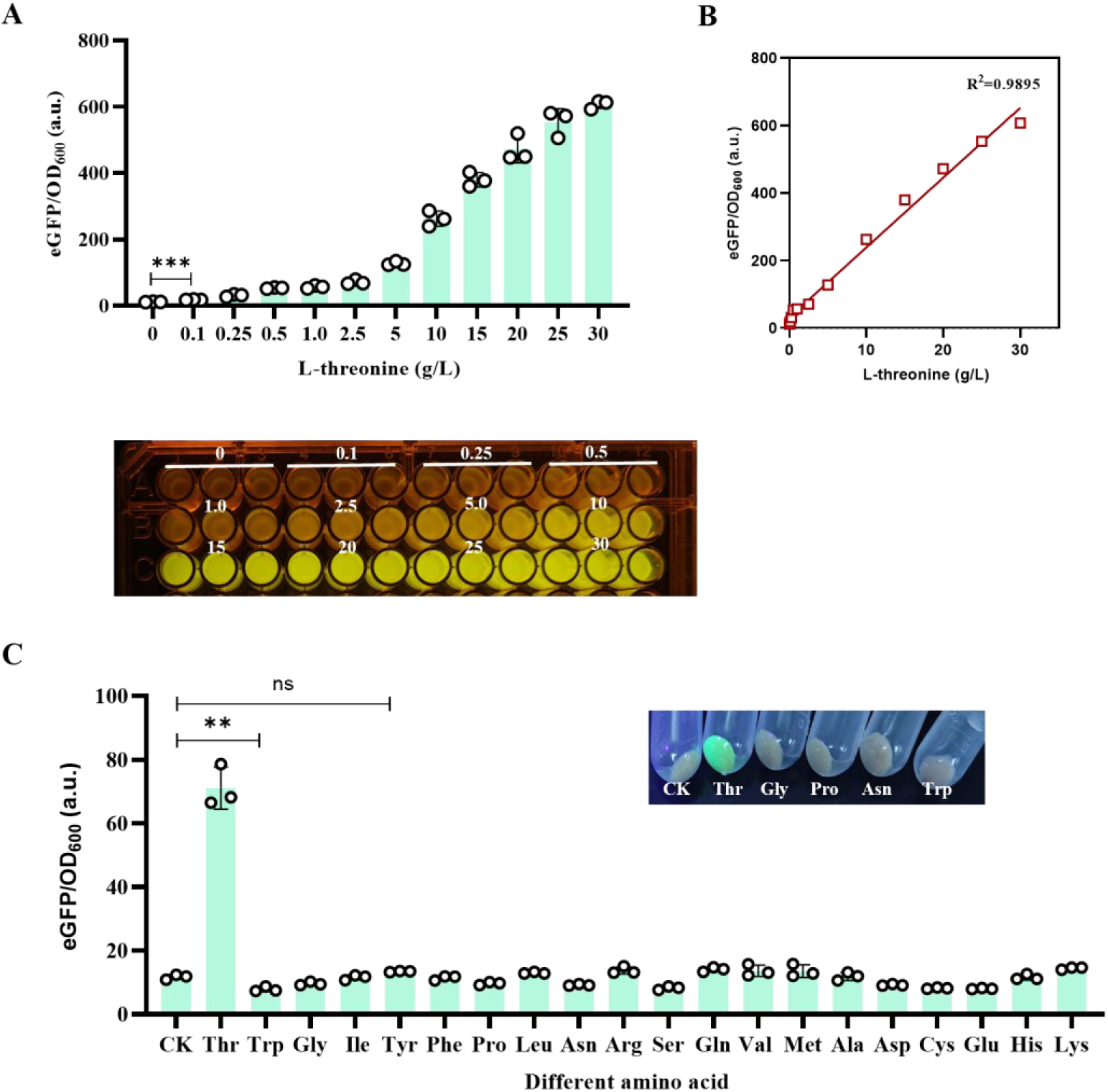
Dose-response and specificity of the optimized –10B biosensor. (A) Response of the –10B biosensor to different concentrations of L-threonine concentrations, with representative well images from a 96-well plate. (B) Recorded response as a function of the applied L-threonine concentration. (C) Response of the –10B biosensor to the 20 amino acids, each at 20 mM, with representative cell images after induction for selected amino acids. ** *p* < 0.01, *** *p* < 0.001, ns, no significant difference by two-tailed Student’s t-test.

A sensor’s practical value depends on its specificity as well as on its dose-response behavior, so we screened a panel of 20 amino acids at a single concentration. Only L-threonine elicited a clear response (Fig. 4C). Tryptophan, the cognate ligand of the Trp repressor, suppressed the reporter fluorescence, consistent with its canonical repressive role in the *trp* system. The other amino acids produced no significant change relative to the control. The sensor was therefore specific for L-threonine across the tested panel, distinguishing it from previously described devices that respond to two or more amino acids. As an additional control, L-threonine from three different commercial suppliers was tested to exclude out an impurity common to particular preparations. The responses were indistinguishable among suppliers (Fig. S1B), confirming that L-threonine itself is the effector.

Existing threonine biosensors tend to compromise on either range or specificity. The *cysJHp* fusion promoter identified through proteomics enabled flow-cytometric screening, but its approximately fourfold signal span provided limited quantitative quantification (Liu et al. 2015). The dual-response circuit based on the *thrL* riboswitch and a *lacI*/P*_trc_* amplification module reached a 4.2-fold dynamic range over 0-20 g/L in our earlier work (Su et al. 2024). Later designs prioritized one property over the other: the CysB-based sensor responds over approximately 0-4 g/L (Zhao et al. 2025b), whereas the SerR^F104I^ regulator recognizes proline as well as threonine (Pu et al. 2025). In comparison, the present sensor operates linearly up to 30 g/L, covering titers typical of shake-flask screening, and its induction ratio exceeds those of the earlier fluorescent reporters by more than an order of magnitude. It also distinguishes L-threonine from all other amino acids tested. This specificity was obtained without engineering the regulator, simply by retaining the operator recognized by the native repressor. The low OFF-state background is central to this performance: because the ratio is measured against a low baseline, it captures the full response rather than a small increment above a high background.

### 3.4 The biosensor discriminates strains with different L-threonine production capacities

A biosensor validated only with the amino acid added to the medium may not respond to the endogenously produced metabolite in the same way, because intracellular concentration, transport, and metabolic state differ from those under supplementation. We therefore tested the –10B sensor in two L-threonine-producing strains with different biosynthetic capacities, measuring both specific fluorescence and the extracellular L-threonine concentration after cultivation. The sensor clearly separated the two strains. The strain with higher L-threonine production produced a correspondingly stronger fluorescence signal, whereas the lower-producing strain produced a weaker signal (Fig. 5B). Across the strains examined, the agreement between the fluorescence readout and the analytically measured titer indicates that the sensor reflects production capacity rather than simply responding to the medium composition.

**Fig. 5.**
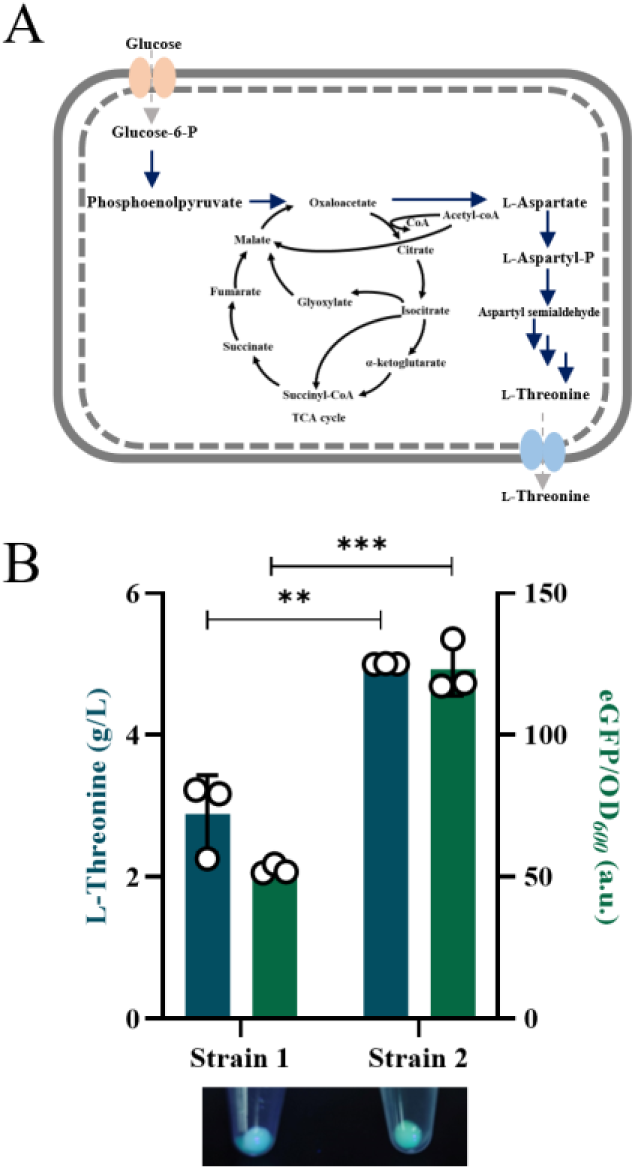
Functional validation in L-threonine-producing strains with different production capacities. (A) Schematic of L-threonine biosynthesis in *E. coli*. (B) Specific fluorescence (eGFP/OD*_600_*) and L-threonine production of two strains with different production capacities, characterized with the –10B biosensor. ** *p* < 0.01, *** *p* < 0.001, significant difference by two-tailed Student’s t-test.

Because the sensor is built from a native *E. coli* regulatory element and the producing strains belong to the same species, it was transferred to the production backgrounds without re-optimization. In strain development, high-throughput screening is often the rate-limiting step, because the phenotypic space generated by random mutagenesis is much larger than what can be assayed by chromatography (Sarnaik et al. 2020; Tuo et al. 2023). A sensor that functions without re-optimization in producing strains is therefore useful for screening mutant libraries by direct measurement.

### 3.5 Biosensor-guided directed evolution of homoserine kinase

As a practical test, the biosensor guided the directed evolution of a key enzyme in L-threonine biosynthesis. Homoserine kinase (ThrB) phosphorylates L-homoserine and is subject to feedback inhibition by L-threonine. In *Corynebacterium glutamicum*, mutations that relieve this inhibition increase threonine accumulation (Petit et al. 2018; Zhao et al. 2025a). The enzyme is highly conserved, and sequence alignment of ThrB orthologs identified residue A12 as a conserved position within the catalytic and regulatory region. We subjected this residue to saturation mutagenesis with an NNK codon.

The mutagenesis design followed our earlier screening workflow (Su et al. 2024), with two deliberate changes to the genetic arrangement. The *thrB* gene was expressed from the pET30a under the P*_J23104_* promoter, whereas the biosensor was carried on the lower-copy pACYC plasmid (Fig. 6A). This arrangement the sensor response within its linear range while permitting high-level expression of the enzyme variants. The mutant library was transformed into WS000 cells already harboring the –10B biosensor, and colonies were screened for increased fluorescence (Fig. 6B). From the library, three mutants (DNA sequencing confirmed) were recovered, and the best variant, designated M4, increased the L-threonine titer by 3.7-fold under the fermentation conditions used (Fig. 6C and D). This supports the use of fluorescence as a sensitive first-pass readout, with analytical quantification reserved for the final candidates.

**Fig. 6.**
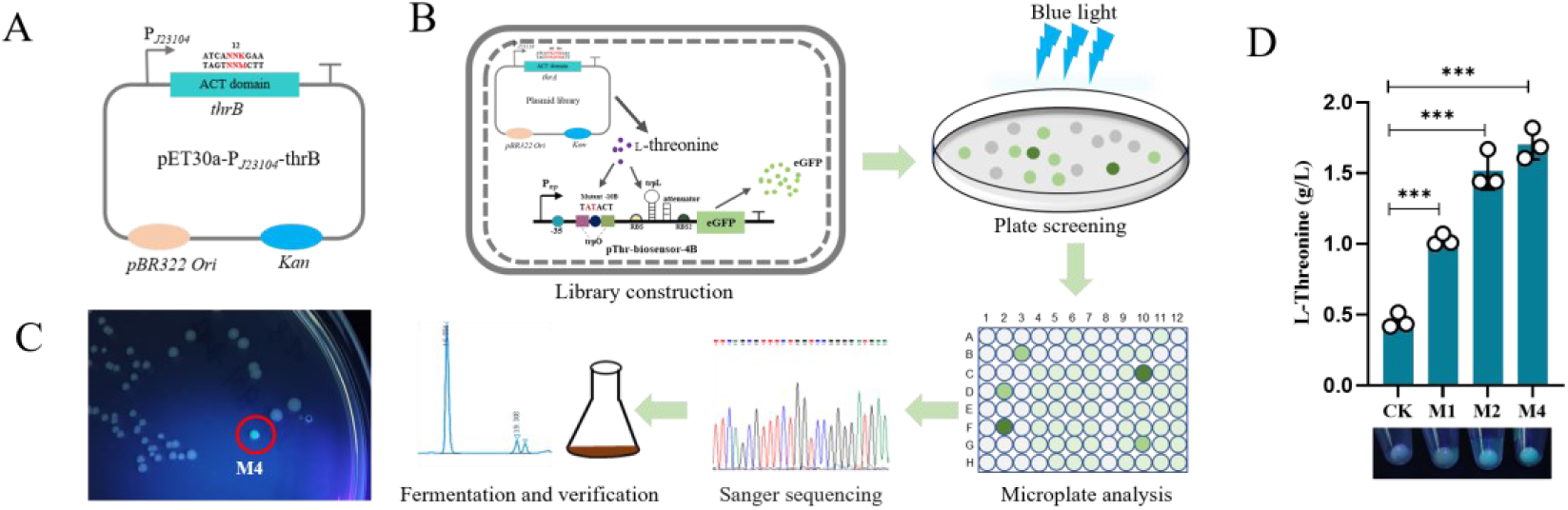
Biosensor-guided directed evolution of homoserine kinase. (A) Schematic of the plasmid carrying the *thrB* gene targeted for mutagenesis. (B) Workflow of saturation mutagenesis: the mutant library was introduced into cells harboring the –10B biosensor: colonies were screened under blue light: elevated green-fluorescent colonies were picked and confirmed by DNA sequencing; validated mutants were cultivated and assayed for fluorescence in 96-well plates; and L-threonine production was confirmed by shake-flask fermentation and HPLC. (C) Mutant strains on the selective plate. (D) L-Threonine production by the mutant strains. *** *p* < 0.001 significant difference by two-tailed Student’s t-test.

The titer gain from ThrB engineering was smaller than the gains reported for the upstream enzyme ThrA, which exerts greater control over pathway flux as the committed step. The native *trp* promoter provided both the specificity and the reporter output, so the sensor required no additional selection device. Alternative screening approaches, such as growth-coupled selection with rare-codon-rich markers (Zheng et al. 2018) or biosensors created by inserting a fluorescent reporter into a ligand-binding domain (Nadler et al. 2016), were therefore unnecessary here. Together with the strain-discrimination results above, these findings establish the biosensor as a practical tool for screening L-threonine producers.

## 4. Conclusions

This work shows that L-threonine induces the tryptophan-repressed *trpEDCBA* promoter of *E. coli*, a response not previously documented. Both the *trpO* operator and the *trpL* attenuation element contribute to the response, and basal expression is lowest only when both elements are present. Stepwise optimization of the ribosome-binding site, the promoter –10 region, the plasmid copy number, and TrpR abundance produced a biosensor that responds linearly to L-threonine from 0.1 to 30 g/L, with a maximum induction of 51.7-fold and specificity for L-threonine among the amino acids tested. The sensor distinguished L-threonine-producing strains with different capacities and supported directed evolution of homoserine kinase, yielding ThrB variants with brighter fluorescence and increased production. The biosensor provides a practical tool based on native *E. coli* regulatory elements for screening L-threonine overproducers. Retaining the native operator of a metabolite-repressed promoter, may also guide the design of sensors for other amino acids.

## CRediT authorship contribution statement

**Buli Su:** Conceptualization, Methodology, Investigation, Data curation, Formal analysis, Visualization, Writing-original draft. **Peixuan Lai:** Investigation. **Xiong Xiong:** Data curation. **Yonghong Wang:** Data curation **Man Li:** Project administration, Writing-review and editing. **Honghui Zhu:** Funding acquisition, Supervision. **Ming-Rong Deng:** Conceptualization, Funding acquisition, Supervision, Writing –review and editing.

## Declaration of generative AI and AI-assisted technologies in the manuscript preparation process

During the preparation of this work the authors used a large language model (DeepSeek) to assist with language editing and to improve the clarity and idiomaticity English of the manuscript. After using this tool, the authors reviewed and edited the content as needed and take full responsibility for the content of the published article. All experimental data, scientific results, and conclusions were generated and verified by the authors.

## Acknowledgements

This work was supported by the GuangDong Basic and Applied Basic Research Foundation (Grant numbers 2026A1515011551) and GDAS’ Project of Science and Technology Development (Grant numbers 2022GDASZH-2022010101).

## Appendix A. Supplementary data

E-supplementary data of this work can be found in online version of the paper.

Table S1 Strains and plasmids used in this study. Table S2. Primers used in this study.

## Data availability

Data will be made available on request.

## Figure captions

The following colour figures are intended for reproduction in colour on the Web and in black-and-white in print.

## References

1. Alabi, T., Adedokun, S., 2025. Amino Acid Nutrition in Poultry: A Review. Animals 15(22), 3323. 10.3390/ani15223323.

2. Ellefson, J.W., Ledbetter, M.P., Ellington, A.D., 2018. Directed evolution of a synthetic phylogeny of programmable Trp repressors. Nat. Chem. Biol. 14(4), 361–367. 10.1038/s41589-018-0006-7.

3. Gao, J., Du, M., Zhao, J., Yue, Z., Xu, N., Du, H., Ju, J., Wei, L., Liu, J., 2022. Design of a genetically encoded biosensor to establish a high-throughput screening platform for L-cysteine overproduction. Metab. Eng. 73, 144–157. 10.1016/j.ymben.2022.07.007.

4. Gong, X., Zhang, R., Wang, J., Yan, Y., 2022. Engineering of a TrpR-Based Biosensor for Altered Dynamic Range and Ligand Preference. ACS Synth Biol 11(6), 2175–2183. 10.1021/acssynbio.2c00134.

5. Hong, K.Q., Zhang, J., Jin, B., Chen, T., Wang, Z.W., 2022. Development and characterization of a glycine biosensor system for fine-tuned metabolic regulation in *Escherichia coli*. Microb Cell Fact. 21(1), 56. 10.1186/s12934-022-01779-4.

6. Hyde, E.I., Ramesh, V., Frederick, R., Roberts, G.C., 1991. NMR studies of the activation of the *Escherichia coli* trp repressor. Eur. J. Biochem. 201(3), 569–579. 10.1111/j.1432-1033.1991.tb16316.x.

7. Lawson, C.L., Sigler, P.B., 1988. The structure of trp pseudorepressor at 1.65 Å shows why indole propionate acts as a trp ‘inducer’. Nature 333(6176), 869–871. 10.1038/333869a0.

8. Li, H., Zhang, W., Han, Y., Tang, G., Lu, F., Qin, H.M., 2025. Programming a bacterial biosensor for directed evolution of tryptophan hydroxylase via high-throughput droplet sorting. Biosens. Bioelectron. 271, 117072. 10.1016/j.bios.2024.117072.

9. Liu, Y., Li, F., Zhang, X., Cao, G., Jiang, W., Sun, Y., Zheng, P., Zhang, D., 2014. A fast and sensitive coupled enzyme assay for the measurement of l-threonine and application to high-throughput screening of threonine-overproducing strains. Enzyme Microb. Technol. 67, 1–7. 10.1016/j.enzmictec.2014.08.008.

10. Liu, Y., Li, Q., Zheng, P., Zhang, Z., Liu, Y., Sun, C., Cao, G., Zhou, W., Wang, X., Zhang, D., Zhang, T., Sun, J., Ma, Y., 2015. Developing a high-throughput screening method for threonine overproduction based on an artificial promoter. Microb Cell Fact. 14, 121. 10.1186/s12934-015-0311-8.

11. Nadler, D.C., Morgan, S.-A., Flamholz, A., Kortright, K.E., Savage, D.F., 2016. Rapid construction of metabolite biosensors using domain-insertion profiling. Nat Commun 7(1), 12266. 10.1038/ncomms12266.

12. Peng, J., Li, X., Li, Y., Zhu, X., Wang, Y., Sun, Z., Zhang, X., 2026. Design and construction of a novel transcription factor-based glycine biosensor for directed enzyme evolution in *Escherichia coli*. Biosens. Bioelectron. 300, 118478. 10.1016/j.bios.2026.118478.

13. Petit, C., Kim, Y., Lee, S.K., Brown, J., Larsen, E., Ronning, D.R., Suh, J.W., Kang, C.M., 2018. Reduction of Feedback Inhibition in Homoserine Kinase (ThrB) of *Corynebacterium glutamicum* Enhances l-Threonine Biosynthesis. ACS omega 3(1), 1178–1186. 10.1021/acsomega.7b01597.

14. Pu, W., Feng, J., Chen, J., Liu, J., Guo, X., Wang, L., Zhao, X., Cai, N., Zhou, W., Wang, Y., Zheng, P., Sun, J., 2025. Engineering of L-threonine and L-proline biosensors by directed evolution of transcriptional regulator SerR and application for high-throughput screening. Bioresour. Bioprocess 12(1), 4. 10.1186/s40643-024-00837-6.

15. Sarnaik, A., Liu, A., Nielsen, D., Varman, A.M., 2020. High-throughput screening for efficient microbial biotechnology. Curr. Opin. Biotechnol. 64, 141–150. 10.1016/j.copbio.2020.02.019.

16. Shi, S., Ang, E.L., Zhao, H., 2018. In vivo biosensors: mechanisms, development, and applications. J. Ind. Microbiol. Biotechnol. 45(7), 491–516. 10.1007/s10295-018-2004-x.

17. Su, B., Lai, P., Deng, M.R., Zhu, H., 2024. Design of a dual-responding genetic circuit for high-throughput identification of L-threonine-overproducing *Escherichia coli*. Bioresour. Technol. 395, 130407. 10.1016/j.biortech.2024.130407.

18. Sun, X., Li, Q., Wang, Y., Zhou, W., Guo, Y., Chen, J., Zheng, P., Sun, J., Ma, Y., 2021. Isoleucyl-tRNA synthetase mutant based whole-cell biosensor for high-throughput selection of isoleucine overproducers. Biosens. Bioelectron. 172, 112783. 10.1016/j.bios.2020.112783.

19. Tuo, J., Nawab, S., Ma, X., Huo, Y.X., 2023. Recent advances in screening amino acid overproducers. Eng Microbiol 3(1), 100066. 10.1016/j.engmic.2022.100066.

20. Wang, Y., Bai, Y., Zeng, Q., Jiang, Z., Liu, Y., Wang, X., Liu, X., Liu, C., Min, W., 2023. Recent advances in the metabolic engineering and physiological opportunities for microbial synthesis of L-aspartic acid family amino acids: A review. Int. J. Biol. Macromol. 253(Pt 3), 126916. 10.1016/j.ijbiomac.2023.126916.

21. Xu, X., Li, X., Liu, Y., Zhu, Y., Li, J., Du, G., Chen, J., Ledesma-Amaro, R., Liu, L., 2020. Pyruvate-responsive genetic circuits for dynamic control of central metabolism. Nat. Chem. Biol. 16(11), 1261–1268. 10.1038/s41589-020-0637-3.

22. Xu, X., Zou, K., Qian, W., Lv, X., Cui, S., Li, Y., Liu, Y., Li, J., Du, G., Chen, J., Liu, L., 2026. Engineering Programmable Tryptophan-Responsive Biosensors Based on RNA-Binding Attenuation Protein for Strain Optimization. ACS Synth Biol 15(2), 610–620. 10.1021/acssynbio.5c00681.

23. Yanofsky, C., 2003. Using studies on tryptophan metabolism to answer basic biological questions. J. Biol. Chem. 278(13), 10859–10878. 10.1074/jbc.X200012200.

24. Yanofsky, C., Platt, T., Crawford, I.P., Nichols, B.P., Christie, G.E., Horowitz, H., VanCleemput, M., Wu, A.M., 1981. The complete nucleotide sequence of the tryptophan operon of *Escherichia coli*. Nucleic Acids Res. 9(24), 6647–6668. 10.1093/nar/9.24.6647.

25. Yu, W., Xu, X., Jin, K., Liu, Y., Li, J., Du, G., Lv, X., Liu, L., 2023. Genetically encoded biosensors for microbial synthetic biology: From conceptual frameworks to practical applications. Biotechnol. Adv. 62, 108077. 10.1016/j.biotechadv.2022.108077.

26. Zhao, G., Tang, Y., Zhang, D., Liu, G., Li, Z., Li, R., Hu, X., Wang, J., Wang, X., 2025a. Engineering homoserine kinase for competitive inhibition release to enhance l-threonine production in *Corynebacterium glutamicum*. Int. J. Biol. Macromol. 321(Pt 4), 146536. 10.1016/j.ijbiomac.2025.146536.

27. Zhao, Z., You, J., Shi, X., Cai, M., Zhu, R., Yang, F., Xu, M., Shao, M., Zhang, R., Zhao, Y., Rao, Z., 2024. Multi-module engineering to guide the development of an efficient L-threonine-producing cell factory. Bioresour. Technol. 416, 131802. 10.1016/j.biortech.2024.131802.

28. Zhao, Z., Zhu, R., Shi, X., Yang, F., Xu, M., Shao, M., Zhang, R., Zhao, Y., You, J., Rao, Z., 2025b. Combining biosensor and metabolic network optimization strategies for enhanced L-threonine production in *Escherichia coli*. Biotechnol Biofuels Bioprod 18(1), 37. 10.1186/s13068-025-02640-7.

29. Zheng, B., Ma, X., Wang, N., Ding, T., Guo, L., Zhang, X., Yang, Y., Li, C., Huo, Y.-X., 2018. Utilization of rare codon-rich markers for screening amino acid overproducers. Nat Commun 9(1), 3616. 10.1038/s41467-018-05830-0.

30. Zhou, G.J., Zhang, F., 2023. Applications and Tuning Strategies for Transcription Factor-Based Metabolite Biosensors. Biosensors 13(4), 428. 10.3390/bios13040428.

