## Supplementary material for "An L-threonine-inducible biosensor derived from the tryptophan-repressed *trp* promoter of *Escherichia coli* for high-throughput screening of L-threonine overproducers": Table S1, Table S2, FIG S1

**Table S1** Strains and plasmids used in this study.

| Strain/plasmid | Description | Source |
|---|---|---|
| *Strains* | | |
| DH5α | *supE44 ΔlacU169 (φ80 lacZΔM15) hsdR17 recA1 endA1 gyrA96 thi-1 relA1* | Lab stock |
| W3110 | Wide type, *λ- F- mcrA mcrB IN (rrnD-rrnE)1* | Lab stock |
| Strian 1 | W3110, *thrL::trc (ΔlacO), ΔptsG, ΔpykF, ΔgltA, ΔpflB, ΔadhE::trc (ΔlacO)-glk-galP, ΔldhA::trc (ΔlacO)-thrA*, Δcrr, ΔlacI, ΔtdcC, ΔiclR::tdcA-gltA-pykF, ΔacrA::grcA-aspC-rhtC-asd-ppc* | Lab stock |
| Strian 2 | W3110, *thrL::trc (ΔlacO), ΔptsG, ΔpykF, ΔgltA, ΔpflB, ΔadhE::trc (ΔlacO)-glk-galP, ΔldhA::trc (ΔlacO)-thrA*, Δcrr, ΔlacI, ΔtdcC, ΔiclR::tdcA-gltA-pykF, ΔacrA::grcA-aspC-rhtC-asd-ppc, ΔpoxB::grcA-aspC-rhtC-asd-ppc, ΔadiA::tdcA-gltA-pykF* | Lab stock |
| WS000 | W3110, *thrL::trc (ΔlacO), ΔptsG, ΔpykF, ΔgltA, ΔpflB, ΔadhE::trc (ΔlacO)-glk-galP, ΔldhA::trc (ΔlacO)-thrA*, Δcrr* | Lab stock |
| Wtrp-ΔtrpR | W3110, *ΔtrpR* | This study |
| *Plasmids* | | |
| pACYCDuet-1 | P15A-derived vector; $Cm^r$ | Lab stock |
| pCDFDuet-1 | CloDF13-derived vector; $Sm^r$ | Lab stock |
| pET30a | pBR322-derived vector; $Km^r$ | Lab stock |
| pSC101 | pSC101-derived vector; $Km^r$ | Lab stock |
| pTargetF-cadA | pMB1 aadA sgRNA-cadA | Lab stock |
| pCas9 | *repA101(Ts) kan Pcas-cas9 ParaB-Red lacIq Ptrc-sgRNA-pMB1*; $Km^r$ | Lab stock |
| pThr-biosensor-1 | pACYCDuet-1 plasmid with *eGFP* under $P_{trp}$ promoter with trpO and trpL, and with original RBS | This study |
| pThr-biosensor-2 | pACYCDuet-1 plasmid with *eGFP* under $P_{trp}$ promoter with trpO and trpL, and with strong RBS | This study |
| pThr-biosensor-3 | pACYCDuet-1 plasmid with *eGFP* under $P_{trp}$ promoter with trpO and without trpL, and with strong RBS | This study |
| pThr-biosensor-4A | pACYCDuet-1 plasmid with *eGFP* under $P_{trp}$ promoter with trpO and trpL, and with strong RBS and mutant pribow box | This study |
| pThr-biosensor-S1 | pACYCDuet-1 plasmid with *eGFP* under $P_{J23119}$ promoter with trpO and trpL, and with strong RBS | This study |
| pThr-biosensor-S2 | pACYCDuet-1 plasmid with *eGFP* under $P_{J23119}$ promoter with trpO and without trpL, and with strong RBS | This study |
| pThr-biosensor-S3 | pACYCDuet-1 plasmid with *eGFP* under $P_{J23119}$ promoter without trpO and trpL, and with strong RBS | This study |
| pThr-biosensor-S4 | pACYCDuet-1 plasmid with *eGFP* under $P_{J23119}$ promoter and with strong RBS | This study |

| | | |
|---|---|---|
| pThr-biosensor-5 | pCDFDuet-1 plasmid with *eGFP* under $P_{trp}$ promoter with trpO and trpL, and with strong RBS and mutant pribow box | This study |
| pThr-biosensor-6 | pET30a plasmid with *eGFP* under $P_{trp}$ promoter with trpO and trpL, and with strong RBS and mutant pribow box | This study |
| pThr-biosensor-7 | pSC101 plasmid with *eGFP* under $P_{trp}$ promoter with trpO and trpL, and with strong RBS and mutant pribow box | This study |
| pThr-biosensor-8 | pACYCDuet-1 plasmid with *eGFP* under $P_{trp}$ promoter with trpO and trpL, and with strong RBS and mutant pribow box; overexpression of *trpR* under $P_{J23119}$ promoter | This study |
| pThr-biosensor-9 | pACYCDuet-1 plasmid with *eGFP* under $P_{trp}$ promoter with trpO and trpL, and with strong RBS and mutant pribow box; overexpression of *trpR* under $P_{J23104}$ promoter | This study |
| pThr-biosensor-10 | pACYCDuet-1 plasmid with *eGFP* under $P_{trp}$ promoter with trpO and trpL, and with strong RBS and mutant pribow box; overexpression of *trpR* under $P_{J23110}$ promoter | This study |
| pThr-biosensor-11 | pACYCDuet-1 plasmid with *eGFP* under $P_{trp}$ promoter with trpO and trpL, and with strong RBS and mutant pribow box; overexpression of *trpR* under $P_{native}$ promoter | This study |
| pThr-biosensor-4B | pACYCDuet-1 plasmid with *eGFP* under $P_{trp}$ promoter with trpO and trpL, and with strong RBS and mutant pribow box | This study |

$Sm^r$, streptomycin; $Cm^r$, chloramphenicol; $Km^r$, kanamycin; R, resistance.

**Table S2.** Primers used in this study.

| Primers | Sequence (5' to 3') |
|---|---|
| **Construction of biosensors** | |
| trpL-F0228 | tcaaatgtagcacctgaagtggctgtgcaggtcgtaaatca |
| trpL-R0228 | gctcctcgcccttgctcaccattgttattctctaattttgttc |
| eGFP-F0228 | gtgagcaagggcgaggagctg |
| pACYC-R1113 | acttcaggtgctacatttgaag |
| RBS-trpL-F0312 | taactttaagaaggagatatacatatggtgagcaagggcgaggag |
| RBS-trpL-R0312 | atatctccttcttaaagttacaaaaaaaagcccgctcattaggc |
| PtrpO-eGFP-F | taactttaagaaggagatatacatatggtga |
| PtrpO-eGFP-R | gtatatctccttcttaaagttagaacttgcgtactagttaacta |
| **Mutation of pribnow box(-10) for Ptrp** | |
| trpL(-10A)-F | cgaactagtataatagtacgcaagttcacgtaaaaag |
| trpL(-10A)-R | gaacttgcgtactattatactagttcgatgattaattgtcaacagc |
| **Deletion of trpR** | |
| trpR-CRISPR-F | ggcatcgcgacgattacgcggttttagagctagaaatagc |
| trpR-CRISPR-R | cgcgtaatcgtcgcgatgccactagtattatacctaggac |
| trpR-UP-F0324 | aggggccagcggcttgatgcagat |
| trpR-UP-R0324 | cctacaaaatcatggagatcgttttggtaggcattc |
| trpR-DOWN-F0324 | acgatctccatgattttgtaggcctgataagacgtgg |
| trpR-DOWN-R0324 | agagcggtgtaccggaacagccct |
| trpR-check-R0324 | atttcacgagatcttcggcga |
| **Construction of biosensors with PJ23119** | |
| pJ23119-trpL-F0305 | ttgacagctagctcagtcctaggtataatgctagcacgtaaaaagggtatcgacaatgaaagc |
| pJ23119-ACYC-R0114 | aggactgagctagctgtcaaacttcaggtgctacatttgaag |
| pJ23119-trpL-F0330 | taggtataatgctagcgaactagttaactagtacacgtaaaaagggtatcgaca |
| pJ23119-R0330 | gtgtactagttaactagttcgctagcattatacctta |
| PtrpO-eGFP-F | taactttaagaaggagatatacatatggtga |
| pJ23119-trpO-(-trpL)eGFP-R0402 | gtatatctccttcttaaagttagtactagttaactagttcgct |
| **Optimization of the plasmid ori** | ggctgtgcaggtcgtaaatca |
| trpL-eGFP-F0401 | caaaaaacccctcaagacccg |
| trpL-eGFP-R0401 | ggtcttgaggggtttttgctgaaaggaggaactatatccg |
| pET30a-trpL-F0401 | atttacgacctgcacagcctgactatcgtcgccgcacttatg |
| pET30a-trpL-R0401 | |
| pCDF-trpL-F0401 | ggtcttgaggggtttttgctgaaacctcaggcatttgaga |
| pCDF-trpL-R0401 | atttacgacctgcacagccgcggttcagtagaaaagatcaaag |
| pSC101-trpL-F0401 | ggtcttgaggggtttttgattgactctcttccgggcgctat |
| pSC101-trpL-R0401 | atttacgacctgcacagcctctttgaatttctccttgattatt |
| **Different overexpression of trpR** | |
| trpR-F0402 | aggtataatgctagctaaggcgctaaggagaccttaaatggcccaacaatcaccctat |
| trpR-R0402 | atttacgacctgcacagccgcgcctgatgcgacgctgccgcg |
| trpL-F0402 | ggctgtgcaggtcgtaaatca |
| pACYC- trpR-R0402 | tagctagcattatacctaggactgagctagctgtcaaacttcaggtgctacatttgaa |

| | |
|---|---|
| PJ23110-trpR-F0409 | caaatgtagcacctgaagttttacggctagctcagtcctaggtacaatgctagctaaggcgcta aggagaccttaa |
| pACYC-R0409 | acttcaggtgctacatttgaa |
| PJ23104-trpR-F0409 | caaatgtagcacctgaagtttgacagctagctcagtcctaggtattgtgctagctaaggcgcta aggagaccttaa |
| Pro-trpR-F0409 | caaatgtagcacctgaagttccgcacgtttatgatatgcta |
| **Mutation of pribnow box(-10) for Ptrp** | |
| trpL(-10B)-F0409 | cgaactagtatactagtacgcaagttcacgtaaaaag |
| trpL(-10B)-R0409 | gtgaacttgcgtactagtatactagttcg |
| **Mutation of thrB** | |
| tdcA-gltA-F | cggcgttacagtaaacgaagcaac |
| tdcA-gltA-R | gcaaaaaacccctcaagacccgtt |
| pACYC-trpL-eGFP-F | gtcttgaggggtttttttgcggctgtgcaggtcgtaaatca |
| pACYC-trpL-eGFP-R | ttcgtttactgtaacgccgacttcaggtgctacatttgaag |
| pET30a-PJ23104-F0712 | gcacgagtactggaaaactaaaataactagcataaccccttggg |
| pET30a-PJ23104-R0712 | gctagcacaatacctaggactgagctagctgtcaatgactatcgtcgccgcactta |
| thrB-F0712 | gtcctaggtattgtgctagctaactttaagaaggagatatacatatggttaaagtttatgccccg |
| thrB-R0712 | ttagttttccagtactcgtgcg |
| thrB-motif-Mu-F | gtttatgccccggcttccagtNNKaatatgagcgtcgggtt |
| thrB-motif-Mu-R0825 | actggaagccggggcataaactttaaccatatgtatatctc |

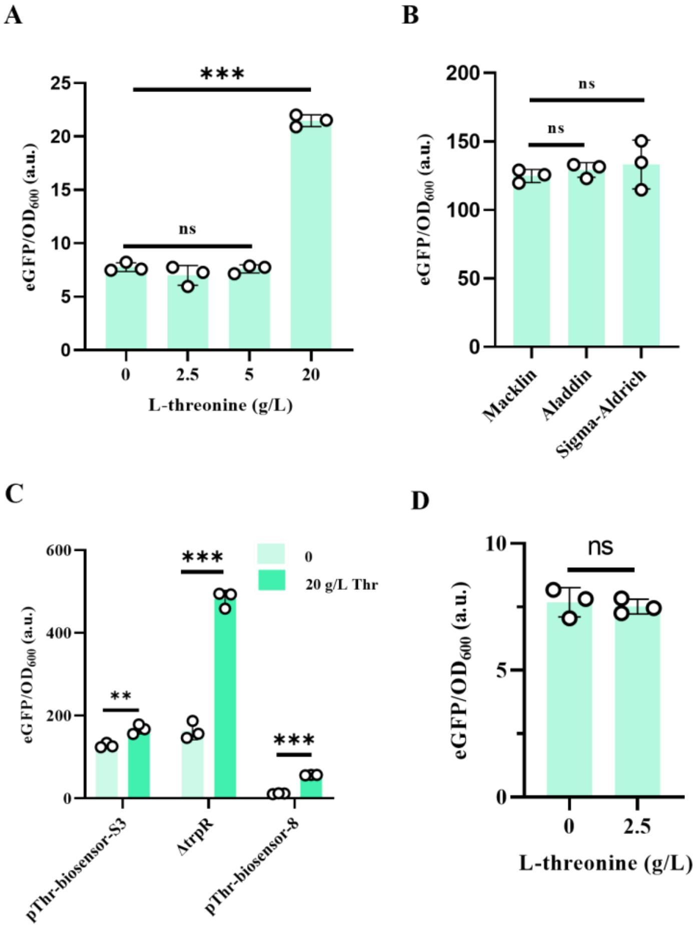


Fig. S1. Supplementary characterization of the biosensor. (A) Specific fluorescence of the -10B sensor with the native RBS (*trpO* and *trpL* present) at different L-threonine concentrations. (B) Response of the biosensor to L-threonine from different commercial suppliers. (C) Response of different sensor constructs to a high concentration of L-threonine. (D) Specific fluorescence of *E. coli* W3110 without the eGFP reporter under different L-threonine conditions, serving as a blank control. ** $p < 0.01$, *** $p < 0.001$, ns, no significant difference by two-tailed Student's t-test.
